# TNF-alpha promotes cilia elongation via Mixed Lineage Kinases signaling

**DOI:** 10.1101/2023.11.30.569502

**Authors:** Amrita Kumari, Amada Caliz, Shashi Kant, Anastassiia Vertii

## Abstract

The primary cilium is characteristic of most of non-immune cells and acts as an environmental signal transduction sensor. The defects in primary cilium have profound consequences on the developmental program, including the maturation of retinal epithelium. The ciliary length is tightly regulated during ciliogenesis. Additionally, many features of ciliogenesis are shared with an immune synapse formation. While the interaction between the cells within an immune synapse is well-characterized, the impact of inflammatory stresses on ciliogenesis in non-immune cells remains elusive. The current study investigates the outcome of inflammatory stimuli for the primary cilium in human retinal epithelial cells. Here, we report that the exposure of retinal epithelium cells to pro-inflammatory cytokine TNF-alpha elongates cilia in a Mixed-Lineage Kinase (MLK) - dependent manner. In contrast, febrile condition-mimicking heat stress dramatically reduced the number of ciliated cells regardless of TNF-alpha exposure, suggesting distinct but rapid effects of inflammatory stresses on ciliogenesis.

## INTRODUCTION

The primary cilium emanates from the basal body (centrosome) in quiescent cells. Upon exiting the cell cycle, the centrosome uses distal appendages on its mother centriole to initiate ciliary vesicle formation and docking to the plasma membrane, a prerequisite for ciliogenesis. The primary cilium serves as an extracellular sensor as it contains many essential for signaling surface receptors such as G-coupled protein receptors and Sonic Hedgehog. Primary cilia also act as a mechanotransduction organelle.

The impact of inflammatory stimuli on cilia is an actively developing area of research. For example, the exposure of cilia to pro-inflammatory cytokine TNF-alpha increased cilia length and was proposed as a specific event for endothelial dysfunction ^1^. Moreover, recruitment of ß-arrestins to the primary cilium via their interaction with intraflagellar transport protein IFT81 and type II bone morphogenetic protein receptor (BMPR-II) within the cilium is essential for endothelial shear stress response ^2^. Inflammatory arthritis is characterized by infiltration of the immune cells in the synovium and increased angiogenesis. A recent report suggests that the regulator of G-protein signaling 12 (RGS12) promotes angiogenesis in inflammatory arthritis via increased ciliogenesis and cilia elongation in endothelial cells, thus suggesting the essential contribution of cilia to endothelial function and angiogenesis ^3^. Another study uncovers the role of ciliary proteins IFT88 in the immune response ^4^, suggesting some ciliary and non-ciliary functions of cilia proteins during inflammation.

Retinal epithelial cells require primary cilium for maturation and development of functional epithelium ^5^. Inflammatory stresses have a detrimental effect on the retinal epithelium. Specifically, inflammation of the retinal epithelium drives the degenerative disorder Mertk-associated retinitis pigmentosa ^6^. We previously reported increased centrosome function in microtubule nucleation during inflammatory stresses in macrophages and retinal epithelial cells ^7^. Additionally, we and others reported fast and detrimental effects of heat stress on centrosome role in ciliogenesis, microtubule organization, and immune synapse formation ^8–13^. Notably, cilia resorption in response to heat stress was reported in mammalian and zebrafish cells ^14^. Given the opposite effects of heat stress and pro-inflammatory cytokines on the centrosome function and the fact that these two inflammatory stresses often co-exist during the immune response and inflammation, we tested the impact of pro-inflammatory cytokine TNF-alpha and febrile-like heat stress alone and in combination on cilia in human retinal epithelial cells.

Within the body, inflammation appears as a complex process involving various components, including inflammatory stresses such as febrile condition and pro-inflammatory cytokines. At the cellular level, the pro-inflammatory cytokine TNF-alpha binds to TNF receptors at the cell membrane and induces a signal transduction cascade, altering the gene expression profile. One of the general mechanisms is the activation of mitogen-activated protein kinases, MAPKs. MAPK kinases p38 and JNK, downstream targets of Mixed Lineage kinases ( MLK1-4), are among the most characterized inflammation-activated stress kinases ^15–17^. Recently, the role of JNK in cilia-related polycystic kidney disease was reported ^18,19^. Therefore, we tested the role of the MLKs-JNK signaling axis in TNF-alpha-induced ciliary changes. Here, we report that in retinal epithelial cells, the exposure to pro-inflammatory cytokine TNF-alpha led to cilia elongation in MLKs-JNK signaling-dependent manner.

## MATERIALS AND METHODS

### Cell culture and treatment

Diploid human retinal pigment epithelium hTERT-RPE 1 cells (Clontech) were grown in DMEM/F-12 (Gibco, #31331-028) supplemented with 10% fetal bovine serum (Gibco, #10500-064), 1% (v/v) Penicillin-Streptomycin (Gibco, #15140-122). For passaging, old medium was aspirated, and cells were washed with 5 ml PBS per 10 cm plate. 1 ml of 0.05% (w/v) trypsin-EDTA was added and the plate was incubated at 37°C until the cells detached. 5 ml RT medium was added per 10 cm plate to quench the trypsin and the cells were transferred to a 15 ml conical tube and pelleted by centrifugation for 5 min at 200g at 4°C. Media was aspirated and cells were resuspended in 5 ml of fresh room temperature warm medium. Cells were passaged at dilutions ranging from 1:2 to 1:10. Cultured human cells between passages 3 to 15 were used for experiments.

Cells treatment regimens are outlined in the table.

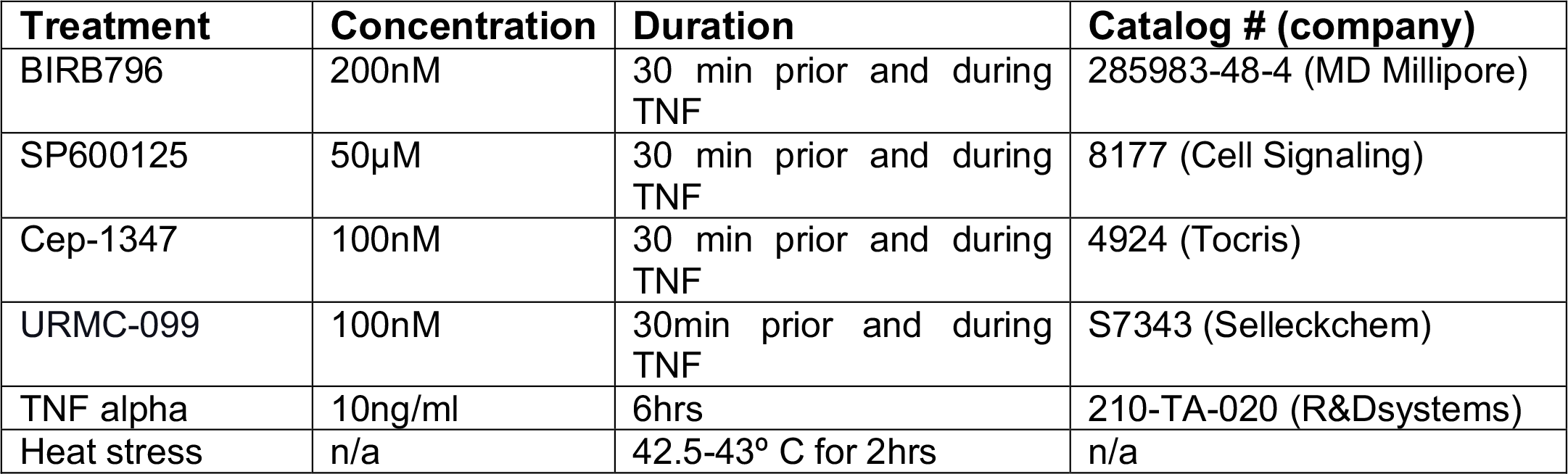

### Immunocytochemistry

Immunocytochemistry was carried as previously described. Briefly, cells were grown on sterile coverslips until ∼70% confluent. After treatment, cells were rinsed with 1xPBS and blocked in 1X PBS/ 1% BSA for 15-30 min at RT. Primary antibodies were diluted in a blocking solution and coverslips were incubated O/N at 4^0^C. Secondary antibodies for immunofluorescence were conjugated with: Alexa 488, Cy3 (Jackson ImmunoResearch, West Grove, PA). Samples were washed with PBS and incubated with secondary antibodies for 1 h at 37 ^0^C. Coverslips were washed with PBS, 1X PBS / 0.1% TritonX-100, and 1xPBS at RT on a shaker, for 10 min each. Finally, the coverslips were rinsed twice with PBS and one time with water, and mounted using Prolong Gold antifade reagent (Invitrogen, cat#P36934).

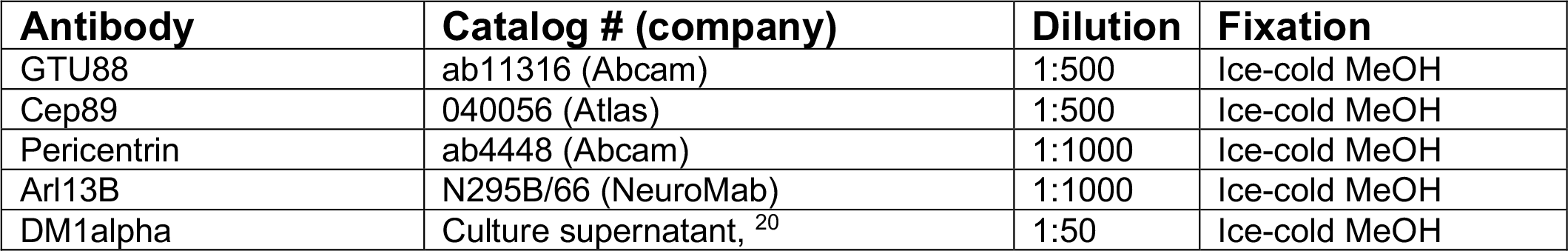

| **Antibody** | **Catalog # (company)** | **Dilution** | **Fixation** |
| --- | --- | --- | --- |
| GTU88 | ab11316 (Abcam) | 1:500 | Ice-cold MeOH |
| Cep89 | 040056 (Atlas) | 1:500 | Ice-cold MeOH |
| Pericentrin | ab4448 (Abcam) | 1:1000 | Ice-cold MeOH |
| Arl13B | N295B/66 (NeuroMab) | 1:1000 | Ice-cold MeOH |
| DM1alpha | Culture supernatant, <sup>20</sup> | 1:50 | Ice-cold MeOH |

### Microscopy, software, and data analysis

Images were acquired with Nikon A1RHD25 (100x NA1.45 Plan-Apocromat Oil objective). Cilia frequencies were counted through Z-stacks manually and were scored as “positive” when adjusted to the number of basal bodies and nuclei throughout the Z-stack. For images, fluorescence range intensity was adjusted identically for each series of panels. *Z* stacks are shown as 2D maximum projections or as a single optical section (Nikon Elements AR software). All images across each experimental series were taken using the same microscope setting (such as laser power and gain) to allow equal comparison of fluorescence levels of samples. All statistical analysis was done using GraphPad Prism software one-way ANOVA test with multiple comparisons or Welch’s *t-*test.

## RESULTS AND DISCUSSION

### The heat stress restricts the number of cells with TNF-alpha-induced cilia elongation

To test the impact of TNF-alpha on cilia length and ciliogenesis, we treated human retinal epithelial cells RPE-1 with either serum starvation media alone or in the presence of TNF-alpha for 24hrs and assessed the frequency of cilia formation and cilia length after 24hrs post-treatment (Figure 1). Our observations suggest that the presence of TNF-alpha in the media induced elongated cilia in comparison to control cells (Figure 1B). These data are in agreement with previously reported effects of TNF-alpha on endothelial cells ^1^. Concomitantly, we detected no significant difference in the frequency of ciliated cells when compared to control and TNF-alpha-exposed cells (Figure 1C).

**Figure 1.**
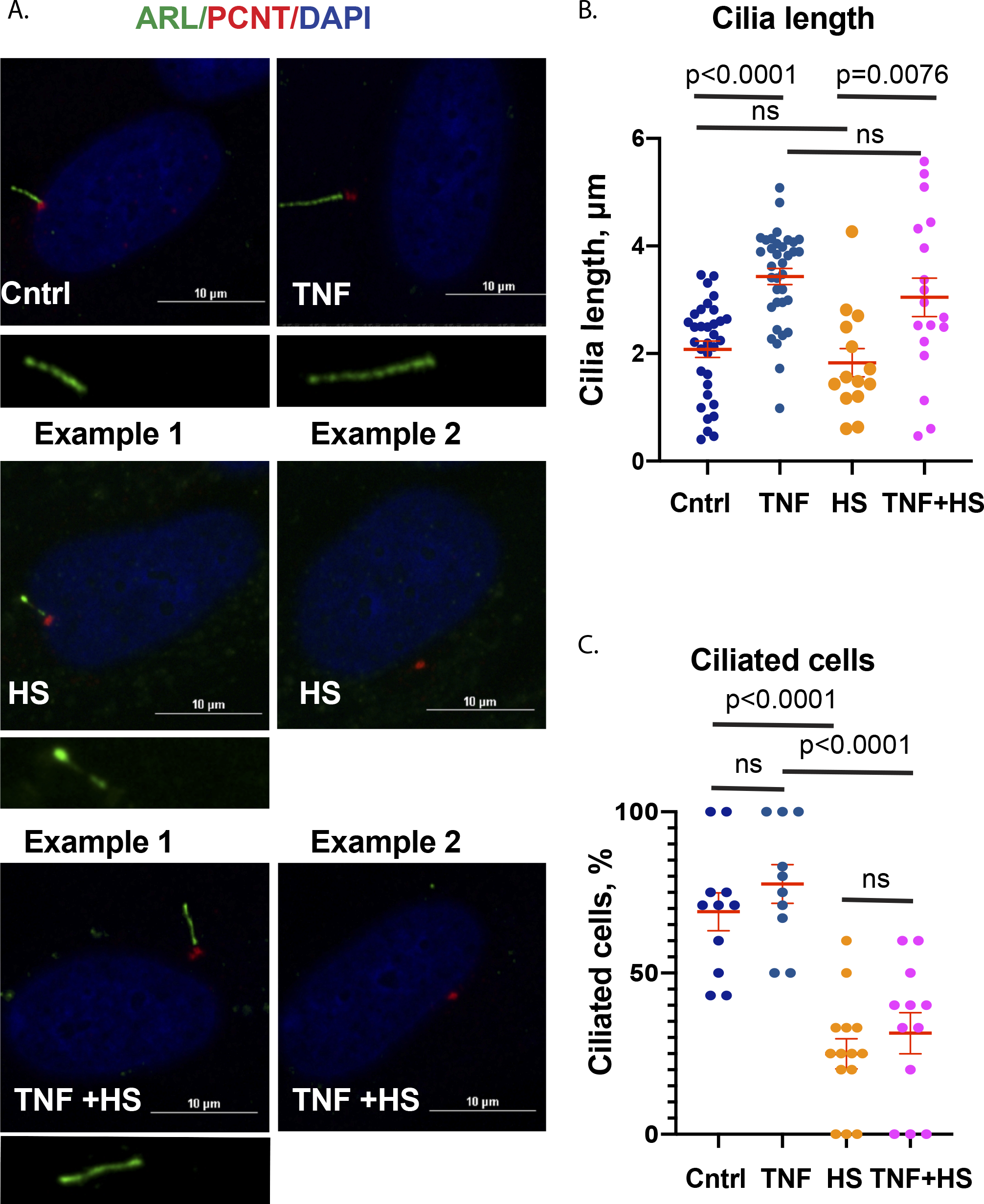
The heat stress restricts the number of cells with TNF-alpha-induced cilia elongation. A. Representative confocal microscopy images of serum-deprived RPE cells that were either left untreated (cntrl), exposed to TNF-alpha for 24 hrs (TNF), heat-stressed for 2hrs (HS), or to a combination of inflammatory stresses (TNF-alpha 22hrs and additional 2hrs under heat stress conditions, TNF+HS). Cilia were visualized using ARL13B antibody (green) and centrosomes (basal bodies) using pericentrin antibody (PCNT in red, scale bar 10µm). B. Cilia length was measured for indicated conditions. For HS–exposed cells, cilia length was measured in ciliated cells (Example 1 from panel A) (n=20-70 cells/sample, micrometers, mean±SEM). C. A percentage of ciliated cells for indicated conditions (a representative graph of a biological triplicate, conditions counted as a percentage of cilia per number of centrosome (basal body) includes Examples 1 and 2 from panel A.

We previously reported that 43ºC heat stress exposure prior to cilia formation led to dramatically decreased ciliogenesis ^8^. Here, we evaluated the effect of the 2-hour-long exposure of HS on the already-formed cilia. Specifically, we treated cells with serum-free media for 22 hours at a usual temperature of 37ºC, and for the final 2 hours, the cells were heated at 43ºC (Figure 1). Although we detected a dramatic decrease in cilia frequency (Figure 1A (example 2 HS) and C), the cilia length of the remaining ciliated cells did not significantly differ from control RPE cells (Figure 1B), possibly arguing against cilia resorption theory as in that case, the gradual decrease in overall ciliary length would be expected prior to the increase in frequency of non-ciliated cells.

A febrile condition is a hallmark of inflammation and appears frequently in combination with pro-inflammatory cytokines such as TNF-alpha. Moreover, TNF-alpha is one of the major pyrogenic cytokines known to induce fever under physiological conditions, including septic shock characterized by uncontrolled febrile condition. To mimic this scenario at the cellular level, we treated RPE cells with a combination of serum-deprived media and TNF-alpha for 22 hours and exposed cells to HS for the next 2 hours. Under this condition, the frequency of ciliated cells was similar to HS alone (Figure A (examples 2 HS and TNF+HS) and C). Of interest, the cilia length of the remaining ciliated cells was comparable to TNF-treated cells and appeared to be elongated in comparison to control cells (Figure 1 A TNF+HS example 1) and B), suggesting that once elongated, cilia are not gradually degraded by consequent treatment with HS. Overall, these data suggest distinct impacts of TNF-alpha and HS on cilia length and frequency. Treatment of HS for 2 hours is sufficient to severely decrease the number of ciliated cells but not the length of the remaining ciliated cells. In contrast, exposure to TNF-alpha does not affect the frequency of cilia formation; however, it induces elongated cilia.

### TNF-alpha induces cilia elongation via MLK signaling

We previously reported the role of Mixed Lineage kinases (MLK2/3) in inflammation-induced centrosome maturation and increased microtubule nucleating capacity in non-ciliated cells ^7^. Therefore, we tested the role of the MLK family in TNF-induced elongated cilia formation. Specifically, we used two small molecule MLK inhibitors, Cep-1347 and URMC-099, for 30 min before the induction of ciliogenesis and during exposure to serum-deprived media with or without TNF-alpha for 24 hours (Figure 2). The presence of MLK inhibitor Cep-1347 had only mild effect on the frequency of ciliated cells during TNF-alpha exposure (Figure 2C). Notably, the elongated cilia length was rescued back to the control length by Cep-1347, and there was a trend in the decreased cilia length in the cells treated with URMC-099 (Figures 2A and B). Thus, our data indicate that TNF-induced activation of MLK signaling is involved in cilia elongation events in RPE cells.

**Figure 2.**
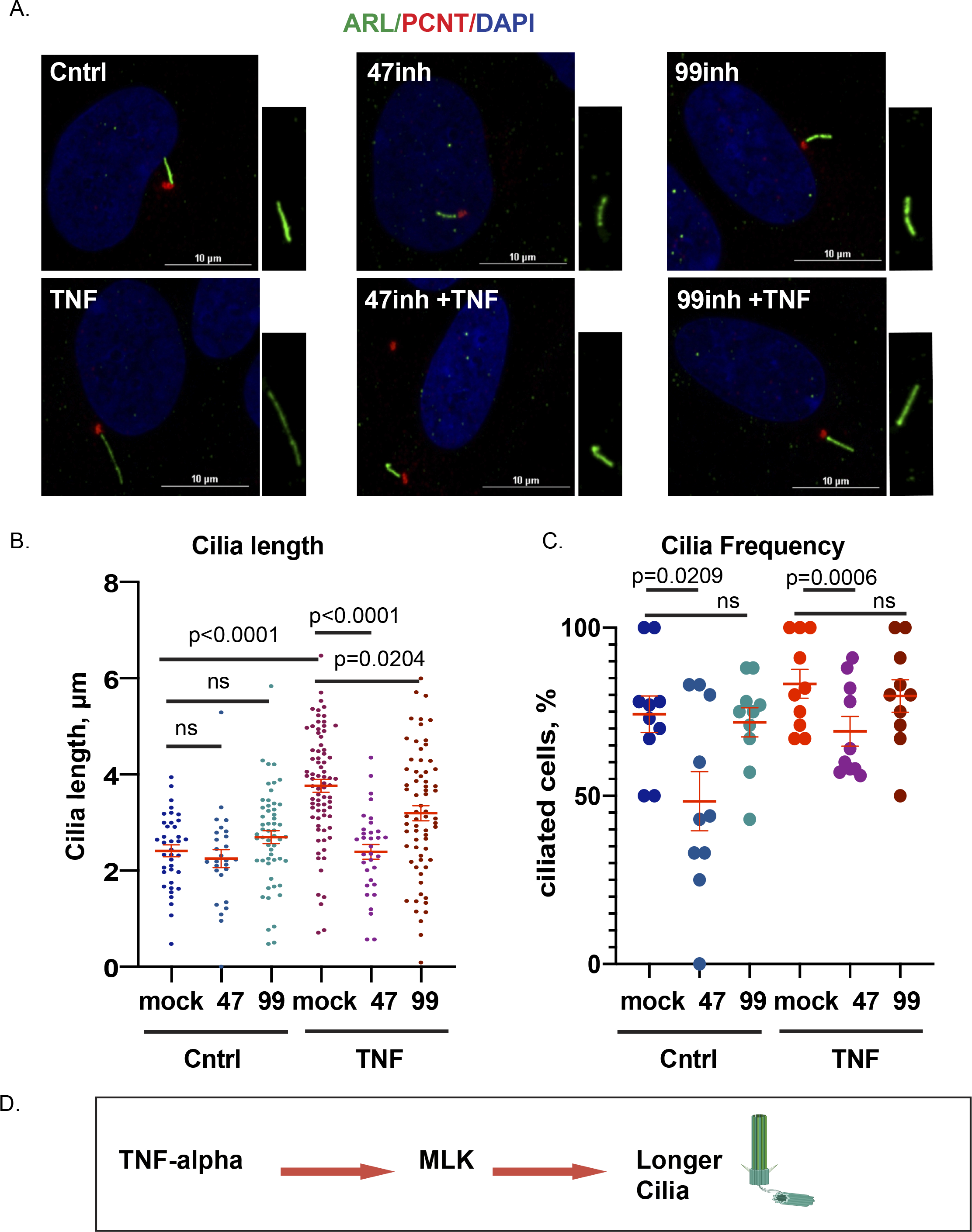
TNF-alpha induces elongated cilia via MLK2/3 signaling. A. Single optical sections of confocal microscopy images of control or exposed to TNF-alpha serum starved RPE cells. When indicated, the cells were pre-treated with MLK2/3 inhibitors Cep-1347 or URMC-099. Cilia were visualized using ARL13B antibody (green) and centrosomes (basal bodies) using pericentrin antibodies (PCNT, red, scale bar 10µm). B. Cilia length was measured for indicated conditions (n=20-70 cells/sample, micrometers, mean±SEM). C. A percentage of ciliated cells for indicated conditions (a representative graph of a biological triplicate, conditions counted as a percentage of cilia per number of centrosomes (basal bodies). D. A model of MLK2/3-mediated cilia elongation.

### JNK but not p38 MAPK contributes to TNF-alpha-induced cilia elongation

TNF-alpha activates multiple downstream MLK targets, with p38 and JNK being among the best-characterized downstream stress kinases ^17^. Moreover, JNK is involved in the cilia-related genetic kidney polycystic disorder ^18,19^ and ciliogenesis in multiciliated pulmonary cells ^21^, and both p38 and JNK were reported to affect atypical centrosome maturation during immune response in non-ciliated macrophages and RPE cells ^7,22^. To evaluate the role of JNK and p38 on TNF-induced cilia elongation and frequency, we used a well-characterized small molecule JNK inhibitor SP-600125 and p38 inhibitor BIRB796 (Figure 3). No dramatic changes in cilia frequency were detected, with slightly lesser frequency for SP600125-treated cells (Figure 3A). The presence of SP600125 but not BIRB796 significantly reduced cilia length (Figure 3B and C), suggesting that JNK but not p38 significantly impacts TNF-induced cilia elongation downstream of MLKs.

**Figure 3.**
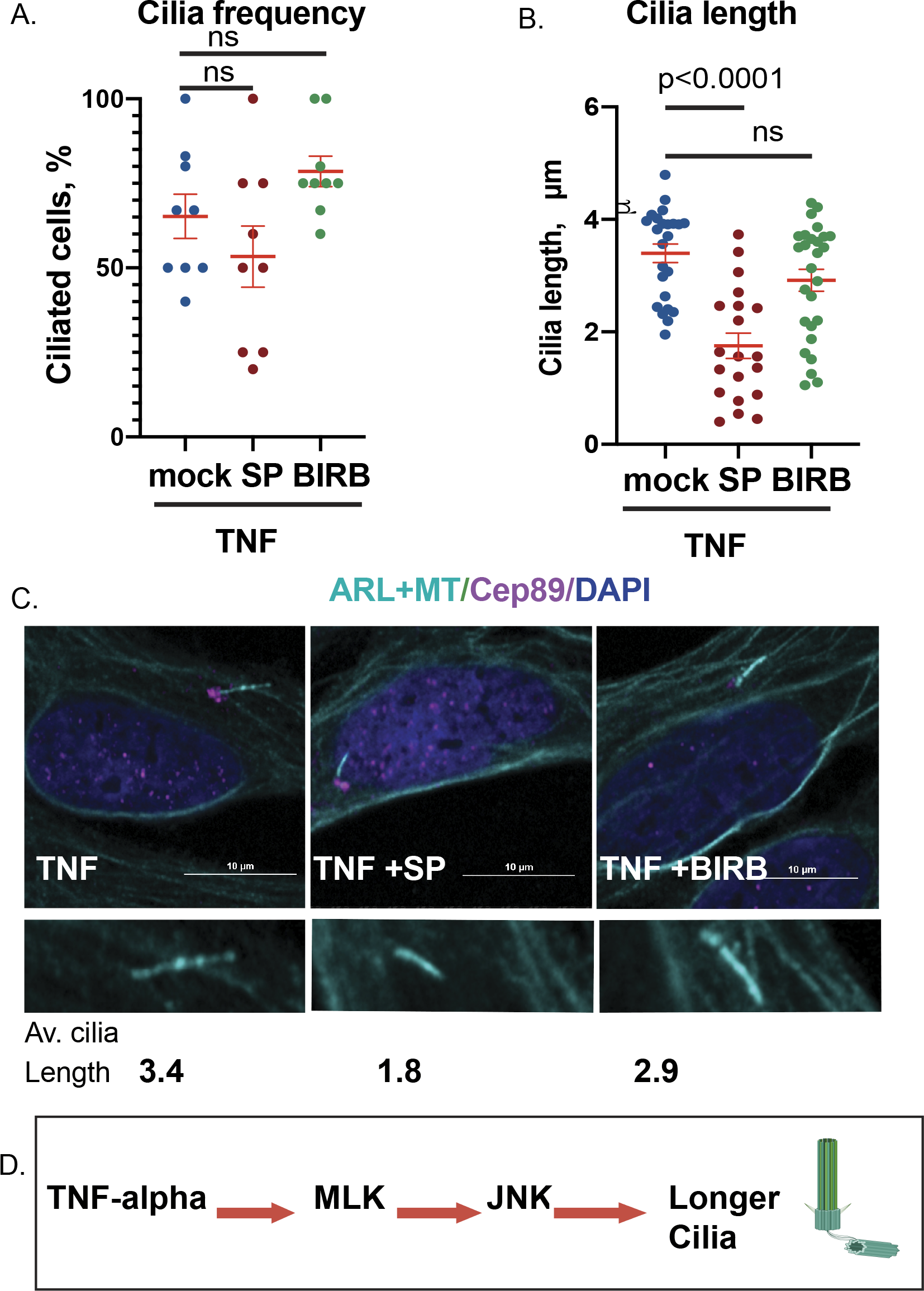
JNK but not p38 MAPK contributes to TNF-alpha -induced cilia elongation. A. A percentage of ciliated cells after serum deprivation in the presence of TNF-alpha either alone or in combination with JNK inhibitor (SP) or p38 small molecule inhibitor (BIRB) (mean±SEM). B. Cilia length was measured for indicated conditions (n=20-70 cells/sample, micrometers, mean±SEM). C. Single optical section confocal microscopy images of cilia and MTs (cyan) and distal appendage, decorated by Cep89 protein (magenta). D. A model of TNF-alpha-induced cilia elongation mediated by MLK-JNK signaling axis.

Inflammatory stresses such as pro-inflammatory cytokines induce febrile conditions and inflammatory responses in immune and non-immune cells. A rapidly expanding research field reveals the impact of inflammation on ciliogenesis and cilia function and the potential role of cilia in inflammatory response and cellular dysfunction ^1,4,23,24^. Our data from human RPE cells are consistent with previous observations from human endothelial cells reporting cilia elongation during exposure to pro-inflammatory ^1,3,25^. Our data also agrees with earlier notion regarding the detrimental effect of heat stress on cilia frequency ^8,14^. We reasoned that pro-inflammatory cytokines have a well-documented pyrogenic effect on the organism, which, at the cellular level, results in a combinatory effect of the two stresses on the ciliogenesis. Therefore, we tested if the combination of the two pro-inflammatory stresses has a different effect on ciliogenesis. Our results suggest no synergetic effect of the two stresses as the addition of febrile-mimicking heat stress does not change the length of the ciliated cell but dramatically decreases the frequency of ciliated cells. Current work also extends the previous observation regarding the potential role of MAPK kinase JNK in ciliogenesis ^18,19,21^, adding the context of the inflammatory response, a previously unreported role for JNK kinase. Generally, p38MAPK and JNK are activated by the MAP kinase kinase kinases (MKKKs or MAP3Ks; such as Mixed Lineage Kinases (MLKs). Mixed lineage kinases are stress-activated protein kinases of the MAP3Ks family ^26, 16^. MLKs possess a characteristic kinase domain that is bordered by an SH3 (SRC homology 3) domain at one end and a leucine zipper domain at the other end, which is followed by a CRIB (Cdc42- and Rac-interactive binding) domain. We have previously shown in systemic MLK2/3-null mice that MLKs are required for the development of insulin resistance and TNF-alpha activates MLKs via the Rho GTPase family members Rac/Cdc42 ^17, 27^. Four family members of MLKs have been identified: MLK1, MLK2, MLK3, and MLK4 ^28^. The expression of MLK1 is primarily seen in the brain, while the other three family members are found widely throughout various tissues ^16,26,28^. Our results point to the MLKs as a new and essential step in the signal transduction cascade that leads to inflammation-induced cilia elongation in human epithelial cells. While function of elongated cilia during the inflammatory response remains unclear, a recent review discussed possible mechanisms and models of cilia elongation and the effects of elongation on cilia function in neuronal cells ^29^. However, the impact of cilia elongation on immune response remains unknown. The elongation appears specific to cytokine treatment as other stresses (such as shear stress in endothelial cells^1^ and heat stress, current work) did not have a similar impact on ciliogenesis.

Overall, we report the previously undocumented role of MLK-JNK signaling in inflammation-induced cilia elongation.

## AUTHORS CONTRIBUTIONS

A.C., A.K. and A.V. made experimental contribution; S.K., provide input and manuscript writing; A.V. developed the concept, made experiments, and wrote the manuscript, and all authors critically read and commented on the manuscript.

## ACKNOWLEDGEMENTS

This work was supported by the American Heart Association Career Development Award to A.V. (grant number 856074). The authors would like to thank Roger Davis, Greg Pazour, Paurav Desai, and Paul Kaufman for support.

## ETHICS APPROVAL STATEMENT

N/A

## PATIENT CONSENT STATEMENT

N/A

